# Linear relationship between peak and season-long abundances in insects

**DOI:** 10.1101/172270

**Authors:** Ksenia S. Onufrieva, Alexey V. Onufriev

## Abstract

1. An accurate quantitative relationship between key characteristics of an insect population, such as season-long and peak abundances, can be very useful in pest management programs. To the best of our knowledge, no such relationship yet has been established.
2. Here we establish a predictive linear relationship between insect catch *M*_*pw*_ during the week of peak abundance, the length of seasonal flight period, *F* (number of weeks) and season-long cumulative catch (abundance) *A* =0.41 *M*_*pw*_*F*. The derivation of the equation is based on several general assumptions, and does not involve fitting to experimental data, which implies generality of the result. A quantitative criterion for the validity of the model is presented.
3. The equation was tested using extensive data collected on captures of male gypsy moths *Lymantria dispar* (L.) (Lepidoptera: Erebidae) in pheromone-baited traps during 15 years.
4. The model was also tested using trap catch data for two species of mosquitoes, *Culex pipiens* (L.) (Diptera: Culicidae) and *Aedes albopictus* (Skuse) (Diptera: Culicidae), in Gravid and BG-sentinel mosquito traps, respectively.
5. The simple, parameter-free equation approximates experimental data points with relative error of 13% and *R*^2^ = 0.997, across all of the species tested.
6. For gypsy moth, we also related season-long and weekly trap catches to the daily trap catches during peak flight.
7. We describe several usage scenarios, in which the derived relationships are employed to help link results of small-scale field studies to the operational pest management programs.

## Introduction

Surveys are crucial for monitoring insect activity, crop pest levels, local movement, long-range migration, feeding and reproduction, and are widely used in pest management programs. Various tools are used for insect surveys, including several types of traps, which are deployed extensively to detect and monitor insect population levels, presence of invading populations and phenological development, for purposes of both applied pest management and research [1-11]. The scale of the enterprise is evident from the following two examples. At least 20 million pheromone lures are produced for monitoring and mass trapping annually worldwide [10]. In the United States, over 100,000 pheromone-baited traps are deployed annually just to monitor gypsy moth [12].

Insect surveys also play an important role in research on population dynamics, seasonal phenology, mating success and mating disruption in various species, including but not limited to gypsy moth, codling moth, light brown apple moth and oriental fruit moth [13-28]. Many of the studies are conducted using releases of laboratory reared insects rather than wild populations, which helps to ensure similar population densities among experimental plots and can provide a longer season of flight for data collection [13, 16-18, 21, 25, 26].

Due to logistics and experimental design, researchers often work with daily or weekly insect catches [17-20, 25-27, 29], while large-scale management programs normally have access to season-long catches only [30], which makes it challenging to relate research results to the management programs To the best of our knowledge, no method currently exists to relate daily or weekly catches to season-long catches. However, relating daily and weekly catches to season-long catches as well as the converse problem of predicting maximum daily catches from the known season-long ones would facilitate interpretation of research results and their application in the management programs. For many species of interest, these population characteristics cannot be assumed constant, which makes it hard to design optimal management protocols. Knowledge of strong correlations between population characteristics can help significantly. Since collecting daily or weekly data in a large-scale management program is very costly and impractical even for a single insect species, a model that relates daily and weekly population characteristics to the easy to obtain season-long catches should be of practical benefit.

More broadly, with about a million insect species currently known [31], it is all but impossible to obtain purely empirical, i.e. based solely on collected data, predictive relationship of the type we seek for even a small fraction of the known species. It may take years or even decades to establish one empirically for a single species, should the need arise. The availability of a theoretical predictive model that is likely to give a quick and reasonable estimate of what to expect should therefore be of value.

## Materials and Methods

### Experimental design and available data

#### Gypsy Moth

We used data from standard USDA milk carton pheromone-baited traps [32] deployed in 2000 and 2001in George Washington National Forest, VA [UTM 637052 E, 4223294 N to 614250 E, 4192715 N, NAD 27, zone 17], in 2001, 2002, 2004 and 2006 in Appomattox-Buckingham and Cumberland State Forests, VA [UTM 746246 E, 4166292 N to 700180 E, 4136389 N, NAD 27, zone 17], in 2013 and 2014 in Goshen Wildlife Management Area, VA [UTM 637052 E, 4223294 N to 614250 E, 4192715 N, NAD 27, zone 17], and in 2016 in Blacksburg, VA [UTM 553841 E, 4121358 N, zone 17]. Permissions to conduct experiments were obtained from VA Department of Forestry and VA Department of Game and Inland Fisheries. At each location, 5 to 20 USDA pheromone-baited traps were deployed, checked and emptied weekly to monitor the flight period of wild gypsy moth populations as part of ongoing research experiments. Captures from all traps in a given location were averaged and considered a single observation. Since 2013, we began to monitor the traps on a daily basis: in 2013, traps were monitored from June 14 to August 4; in 2014, traps were monitored from June 30 to August 8. In both 2013 and 2014, 2 sentinel traps were deployed and checked daily for 5 days each week. In 2016, a trap was placed in Blacksburg, VA and monitored every day from June 20 to August 10.

A standard assumption often made by others [8, 33, 34] is that the distribution of trap captures as a function of time is Gaussian. We have verified this assumption explicitly for two randomly selected years (2006 and 2014) of gypsy moth trap catches, (Fig 1).

**Fig 1:**
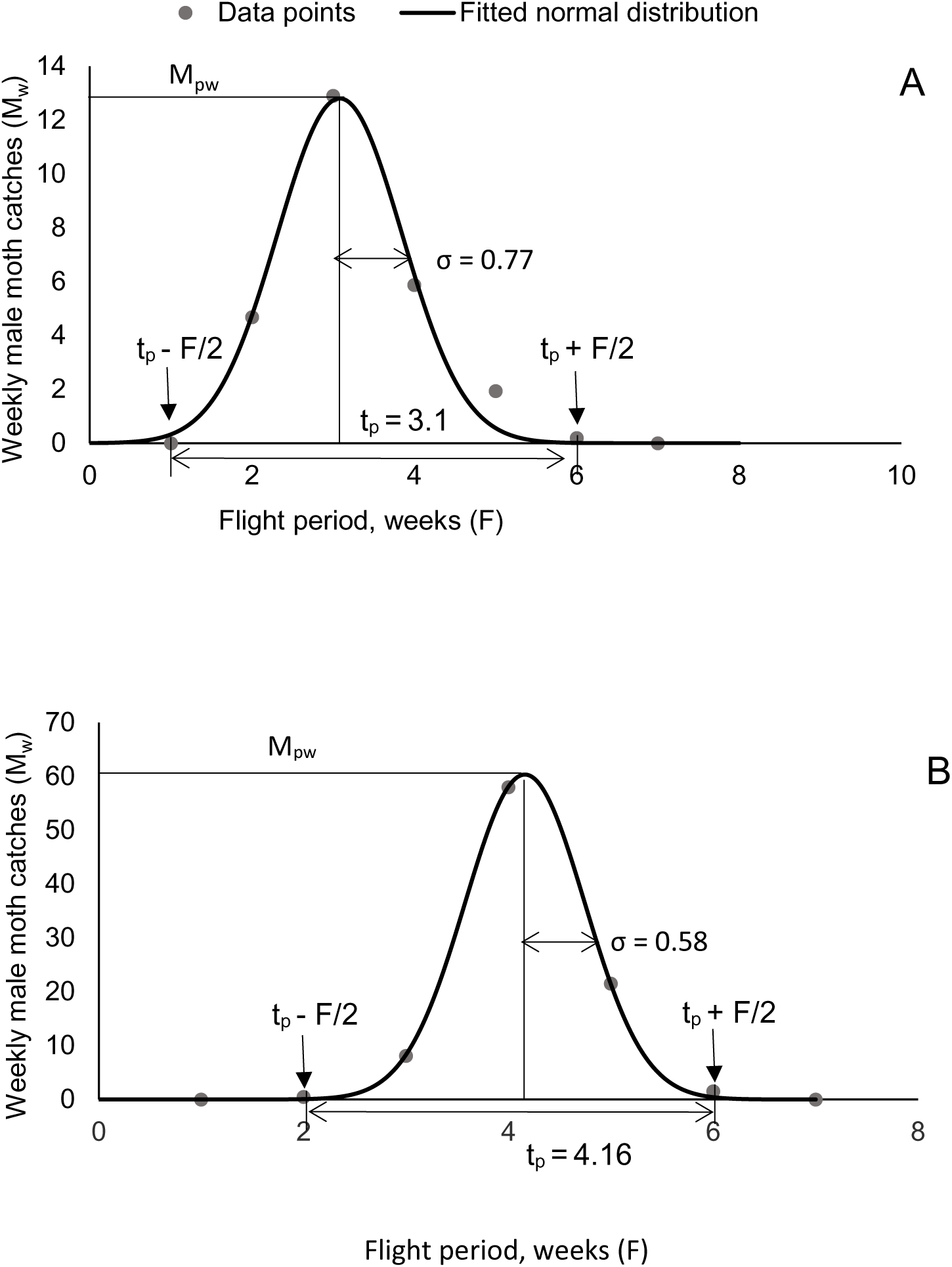
Weekly male gypsy moth catches in pheromone baited traps, 2006 (A) and 2014 (B) and least-square fit of experimental data points to a normal distribution. In both cases, the correlation *R*^2^ > 0.99. Grey arrows indicate the time points where the insect population reaches the trap sensitivity threshold.

#### Mosquitoes

Testing of the proposed model requires high temporal resolution data on insect abundance not readily available for the vast majority of insect species. Fortunately, mosquitoes is an exception. Since in the US, mosquitoes vector important diseases of humans, such as West Nile Virus and Eastern Equine Encephalitis, mosquito risk is often evaluated throughout the season as part of Integrated Mosquito Management Program. Many Mosquito Management Programs conduct weekly trap counts, which are labor intensive and, therefore, are not justifiable for many other pest management programs. We obtained mosquito trapping data from the Mosquito and Forest Pest Management program, Prince William County, VA in 2015. We randomly chose *Culex pipiens* catches in CDC Gravid Traps [35] and *Aedes albopictus* catches in BG-Sentinel traps [36]. The trap data were collected weekly from May 3 to October 23, 2015.

### The main model and its derivation

The model describes the relationship between insect catches *M*_*pw*_ during the week of peak activity, the flight period *F*, and season-long cumulative catches A, which is a direct measure of the insect abundance. It is known that pheromone traps achieve detection probability of nearly 100% even for relatively sparse populations [37]. It is therefore reasonable to assume high sensitivity of the traps. Specifically, we assume that traps are sensitive enough to start catching insects when their population reaches 1% of the maximum population, consistent with our own experiments (see Fig 1 in the results section). We also demonstrate below that the specific value of “trap sensitivity” is not very important for as long as it is high enough – the mathematical structure of the resulting equation is such that it is insensitive to the value of the “sensitivity threshold”. For a symmetric distribution, the flight period F is defined as follows: the traps begin to catch insects at t = t_p_ – F/2, when the insect density rises above the 1% trap sensitivity threshold, and stop at t = t_p_ + F/2, once the insect density drops below this threshold (Fig 2). We model the distribution of insects caught as a function of time as a Gaussian function *M*_*pw*_*e*^-(*t*-*t*^*p*^)^2^/2*σ*^, centered around the peak flight time point *t*_*p*_ (Fig 2). Then, the cumulative catch (A)

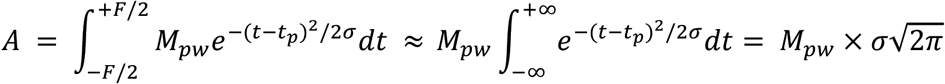

The full width *F* of a Gaussian function at 1/100 (1%) of the maximum is related to σ via

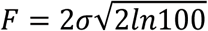

Therefore,

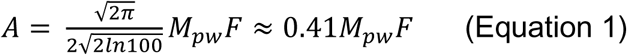

which is the main result of this work. Its robustness to the assumptions made in the derivation is discussed below, and is demonstrated experimentally.

**Figure 2:**
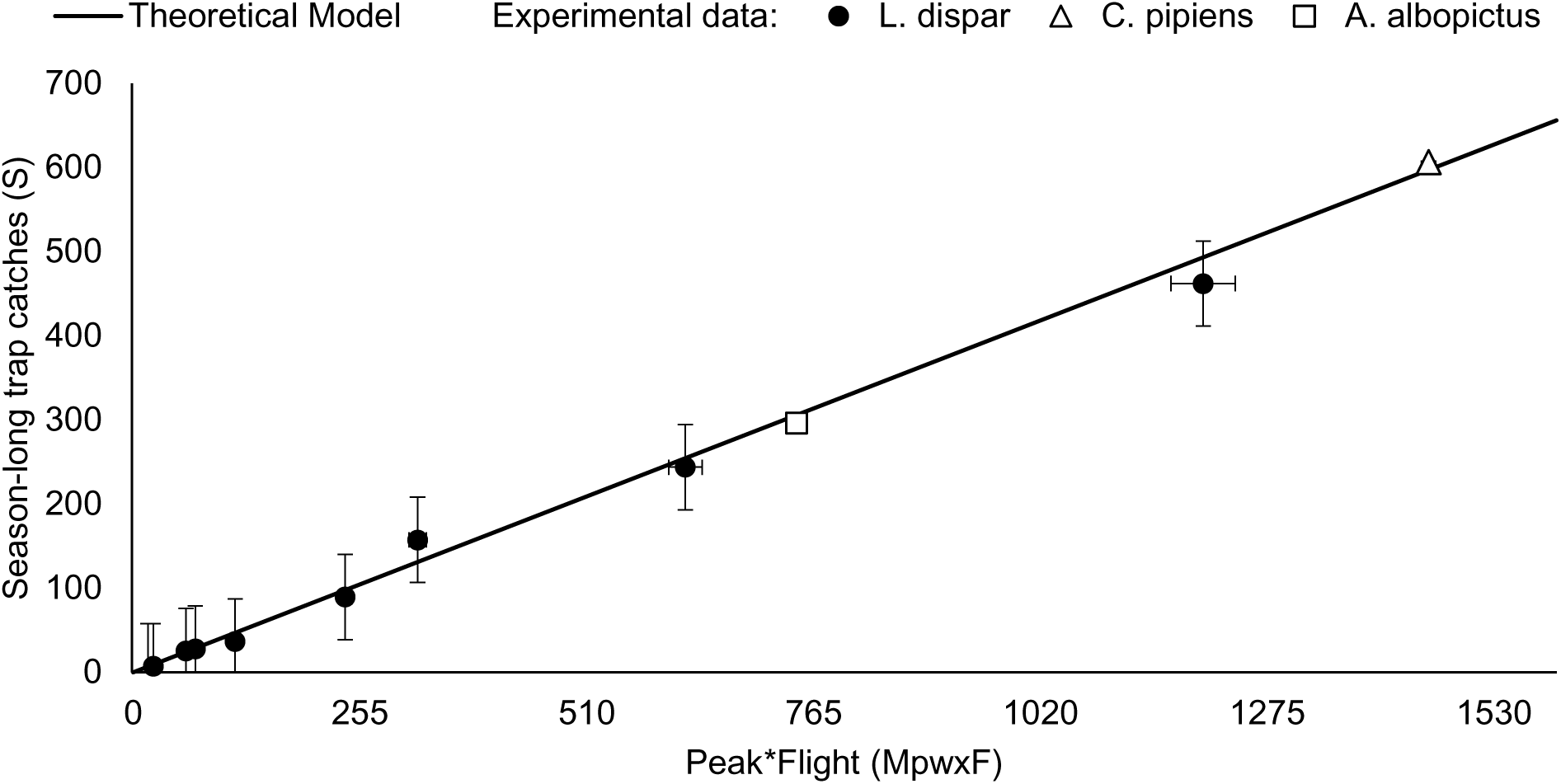
Season-long trap catches depend linearly on the product of trap catch during the peak flight week and the length of the flight period. The relationship, derived on general arguments rather than fitting to experimental data, holds true for several different insect species and catch methods. Error bars (for Gypsy moth): vertical bars represent standard error; horizontal error bards correspond to the temporal resolution of the experimental data points, which is about 3% for most points.

We stress that the derivation of Eq. 1 does not involve any fitting to experimental data. The relationship is expected to hold as long as the following conditions are satisfied: (1) The seasonal change in the insect population as a function of time (weekly averages) can be reasonably approximated by a Gaussian function; (2) The trap method is sensitive enough to detect insects during most of the flight period; (3) Traps do not saturate. The importance of (1) and (2) for our model is self-evident. To understand the importance of the last assumption, consider a situation where it breaks down completely: at very high population density during several weeks, high enough so the traps are completely saturated. Obviously, if this were the case, the weekly catch estimates would be flat around the peak, no longer representative of the true insect abundance [38, 39], and the model that assumes a Gaussian distribution would fail.

### Robustness of the model

Assumptions (2) and (3) essentially state that the catch method is, in some sense, “well designed”. As long as this is the case, Equation 1 is not very sensitive to details such as the exact value of the trap sensitivity threshold. Indeed, consider a trap that faithfully represents the true population density in the following sense: catch = α*(true population density), where the constant α stays the same within a given year and experiment, but may vary for different species and years. Now suppose a different trap is used, which changes the value of α by a factor of 2 for a different species/year. Both *A* and *M*_*pw*_ in Equation 1 will change by a factor of 2, without affecting the linear nature of the relationship. There will be a slight effect on the proportionality constant *K* in *A* = *K*\**M*_*pw*_\**F* due to the use of more/less sensitive trap (e.g. one that can sense 1/200 instead of 1/100 of the maximum population, which will affect the trap sensitivity threshold, Figure 2), but note that the functional dependence of K on the trap sensitivity threshold is extremely weak, 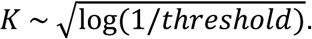 In the above example of the threshold changing from 1/100 to 1/200, the difference in the proportionality constant in Equation 1 is 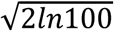 vs. 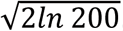, or 3.03 vs. 3.25, which is small.

Assumption (1) has been explicitly verified on our experimental data points for gypsy moth. As we shall see below, even with a relaxed assumption that the seasonal distribution of insect abundance is not strictly Gaussian, but nevertheless has a distinct start, end, and sharp peak, the model is likely to work reasonably well. While assumption (1) (or its relaxed version) is likely to hold for many other species, it implies that the insects are short-lived relative to their entire developmental periods. Otherwise, even if the developmental period itself is normally distributed (which is approximately true even for humans [40]), the abundance can reach a long plateau during the season, inconsistent with the Gaussian shape. For such insects we expect appreciable deviations from the model predictions. For example, tropical species may not have a distinct start and end of flight points, and so the model may not be applicable.

However, as we shall see below, the simple and parameter-free model described by Equation 1 has so far worked surprisingly well for completely different insect species and trapping methods in the temperate climate zones. We stress that we have not used any of the gypsy moth specific parameters in the derivation of Equation 1.

### Relationship between season-long abundance and daily trap catch for gypsy moth

Our main result, Equation 1, describes weekly trap catches. Here we seek to establish a predictive relationship for the daily catches. Unlike the weekly catches, in which day-to-day fluctuations of insect catches are automatically averaged out to produce a smooth distribution that deviates little from the expected Gaussian distribution (Fig 1), individual daily values are too variable for a single characteristic such as daily average to be useful in practice. Here we seek to estimate a conservative range for the upper and lower bounds on the daily trap catches. It is probably hopeless to try to derive the range from first principles; instead, we deduce it from our experimental data for gypsy moth (described below), which we re-interpret as follows. Since the beginning of peak flight week assignment is arbitrary and depends on the day a trap was checked, we simulated different week assignments using a sliding window of 7 days to determine the week of peak flight, making sure that the maximum observed value of daily catch is always included in the “peak flight” week. In effect, the procedure simulates 7 different experimental outcomes from a single data set. The analysis yields a range of proportionality coefficients between trap catch during the peak flight week and the peak daily value.

Daily gypsy moth male trap catches were not available to us prior to 2013. Once daily trap catches became available in 2013, followed by 2014 and 2016, we also sought to relate maximum daily trap catch at peak flight to the season-long trap catch. For this relationship, we used daily trap catches collected in 2014 as a training set, and daily trap catches collected in 2013 and 2016 served as test sets.

In an idealized scenario, without random day-to-day variation, trap catches during the peak week of flight period can be calculated as:

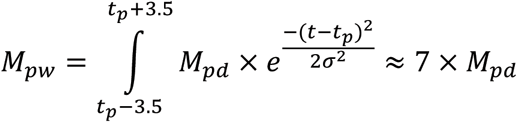

where *M*_*pd*_ is the daily peak, that is the absolute maximum of the distribution. In the idealized case this simple relationship holds because for the species discussed here, *σ* is considerably larger than 7 days, and so the integrand varies little over the integration range in the above expression. However, in reality, daily values fluctuate significantly around the predicted Gaussian peak, resulting in two effects. First, the proportionality coefficient *K* in *M*_*pw*_ = *KM*_*pd*_ becomes less than 7, with daily peak values that deviate stronger from the Gaussian-based expectation resulting in lower *K*. Second, *K* will vary from year to year. In 2014, *K* ranged from 2.35 to 4.66. Below we estimate maximum and minimum bounds on *K*, based on the variation in daily trap catches inferred from our analysis of the experimental data on gypsy moth from 2014. Our estimate is that 2.35*M*_*pd*_ < *M*_*pw*_ < 7*M*_*pd*_, or conversely

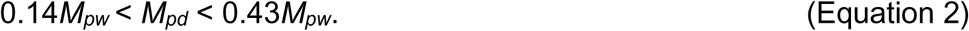

Below we use data points from 2013 and 2016 to validate the estimate.

Using our main result (Equation 1) *A* ≈ 0.41*M*_*pw*_ *F*, we arrive at another useful relationship:

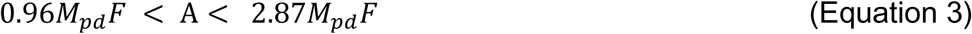

Equations 2 and 3 establish a connection between daily peak values *M*_*pd*_, and other measurable parameters of insect abundance for *gypsy moth*. While the values of the numerical factors in Equation 2 may be species specific, the over-all approach to deriving the inequality is general.

## Results

### Season-long trap catch for gypsy moth

Experimentally observed season-long trap catches are highly variable, showing strong dependence of the season-long cumulative catch (abundance) *A* on both the length of the flight period *F* and the trap catch during the week of peak flight, *M*_*pw*_ (Fig 2). Trap catches during peak flight cannot be predicted from the flight period alone.

### Experimental verification of the model

Our data strongly confirms the proposed functional dependence, Equation 1, between the season-long trap catch, the flight duration and the trap catch during peak flight (Fig 2).

The predicted season-long trap catches from trap catches during the week of peak flight correlate strongly with the actual, measured season-long trap catches for all three species tested (*R*^2^ = 0.9977, SD = 0.13, Fig 2).

We stress the significant range spanned by the data points in our test sets, see “Methods”. The season-long male gypsy moth trap catches span almost two orders of magnitude, ranging from 7.3 to 462 males/trap/week; flight duration ranged from 3.7 to 7 weeks; and maximum trap catch during peak week ranged from 4.4 to 172 males/trap. Season-long trap catches of *C. pipiens* and *A. albopictus* span an order of magnitude, ranging from 6.22 to 60.72 and from 0.12 to 31.12, respectively. Flight duration of both species was 24 weeks.

### Robustness of the model to insect and trap type

Our main result (Equation 1) is robust to insect and trap type (Fig 2).

It is remarkable that the accuracy of Equation 1 is the same for all of the species tested, even though the biology, catch methods, and the geographical areas are different. Moreover, for the mosquito species the catch distributions (Fig. 3) are not as close to idealized Gaussian as they are for the gypsy moth (Fig 1). This observation suggests strong robustness of the proposed model, and its potential to work for other species as well. We explain this robustness as follows, based on the mosquito example. Let’s approximate the actual catch distribution by a triangular shape with the top vertex at the experimental peak point, Fig. 4, rather than by a pure Gaussian. Then, the total catch (abundance) is the area under the triangle *A* = 0.5 × *M*_*pw*_ × *F*. The coefficient 0.5 in this equation is not that different from 0.41 in our main Equation 1. As one can see from Fig. 4, the experimental data points lie mostly below the triangle, which explains why the original formula *A* = 0.41*M*_*pw*_*F* works even better than the one based on the triangle.

**Figure 3:**
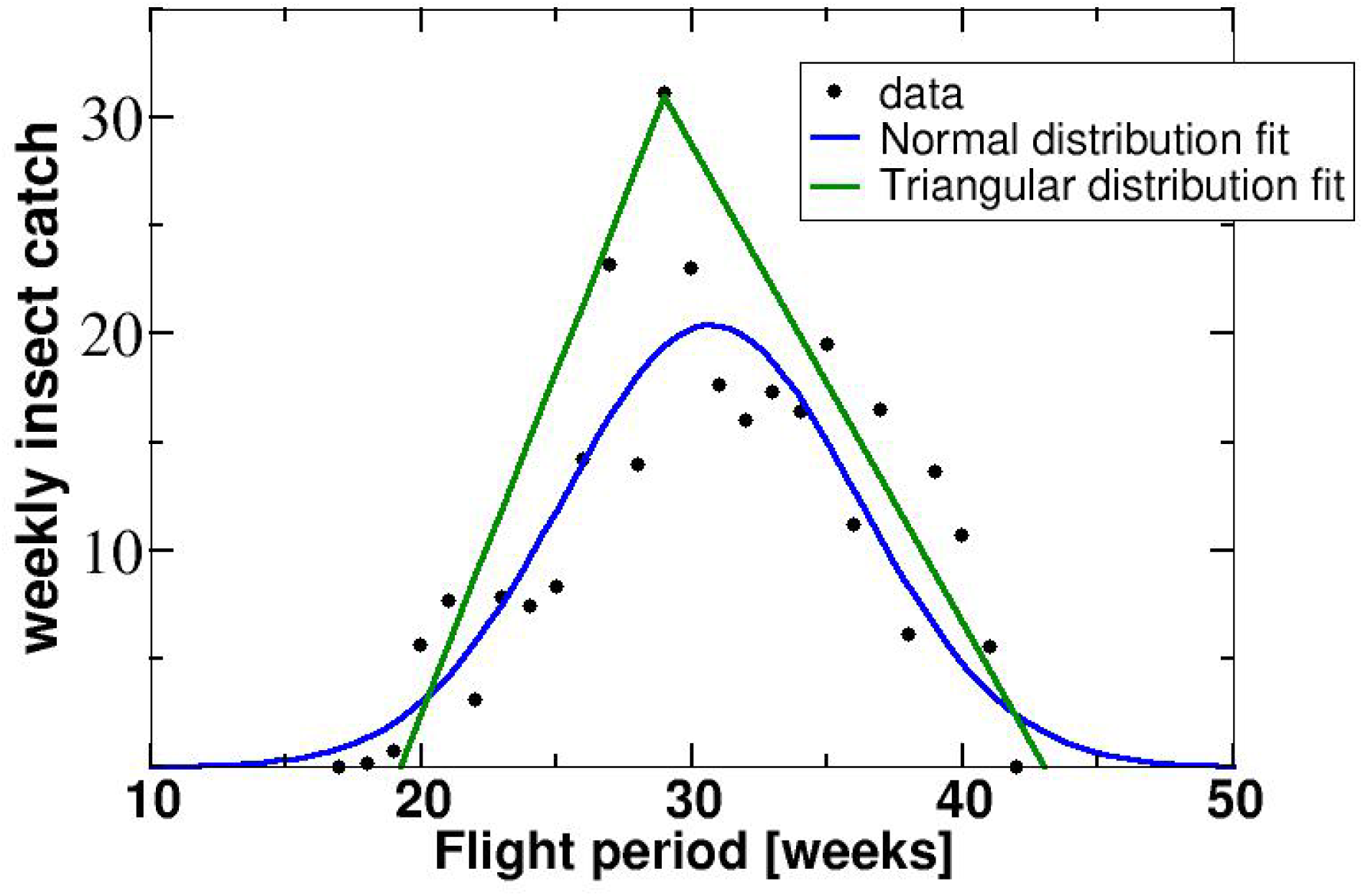
Weekly *A. albopictus* catches in BG-sentinel mosquito traps, 2006 and least-square fits of experimental data points to a normal and a triangular distributions.

To conclude this discussion, consider a hypothetical insect for which Equation 1 does not work because the main assumption on the catch distribution is violated. Namely, consider a catch distribution where instead of a single clear peak there is a long plateau of near constant abundance M_pw_, whose duration P is comparable to the total flight time F. The distribution will look like a trapezoid in this case, giving for the total abundance *A* = 0.5 × *M*_*pw*_ × *F* + 0.5 × *M*_*pw*_ × *P*, where the second term can be considered the error term relative to the main expression *A* = 0.5 × *M*_*pw*_ × *F*. The relative error is then simply P/F, which gives us a simple practical criterion for the application of Equation 1: the time interval over which the population is close to its peak value must be much less than the total flight period.

### Daily trap catch for gypsy moth

Experimental daily trap catches ranged from 0 to 13.5 males/trap/day in 2013, from 0 to 10 males/trap/day in 2014, and from 0 to 15 in 2016. (Fig 4). Clearly, the maximum daily catch during the week of peak flight is within the estimated lower and upper bounds given by Equation 2.

**Figure 4:**
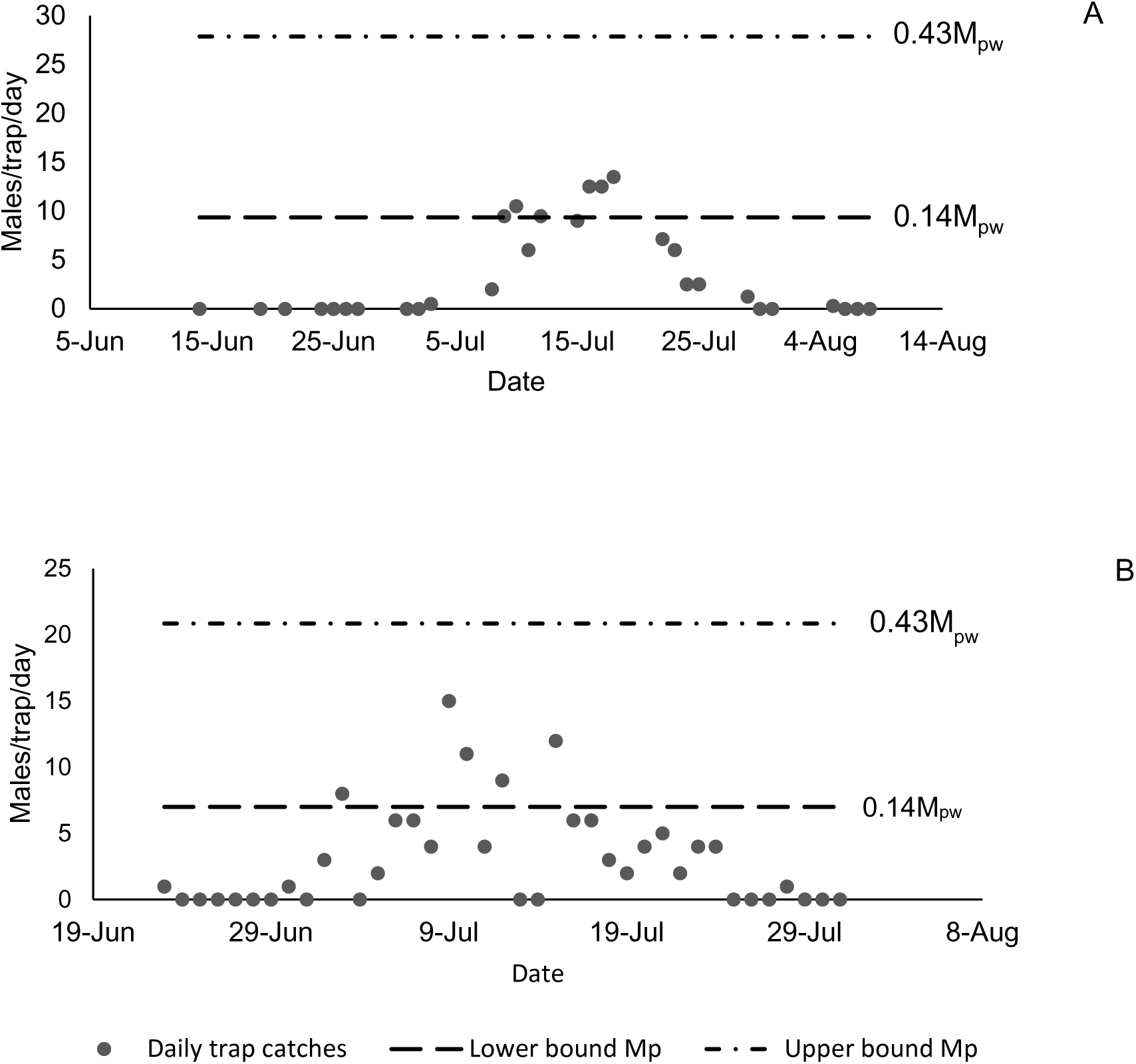
Daily gypsy moth male catches in pheromone-baited traps in Virginia, 2013 (A) and 2016 (B). Lines indicate estimated minimum and maximum possible daily values estimated from the maximum observed weekly peak value.

Equation 3 is tested using the same two data sets. In both years, observec abundance was within the predicted range given by Equation 3 (Table 1).

**Table 1:**
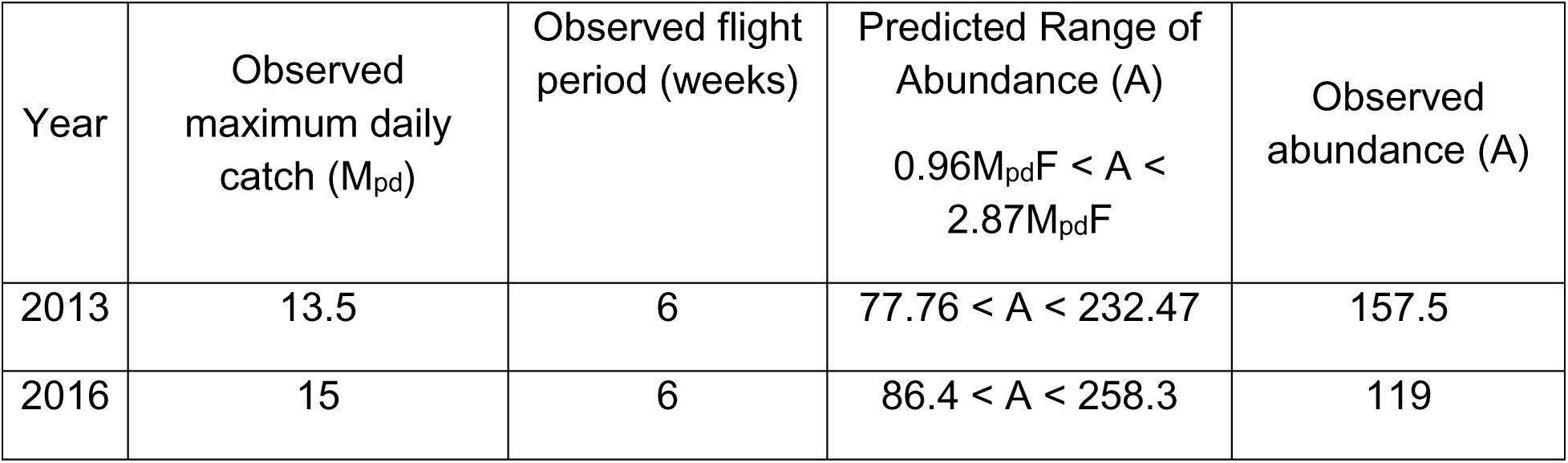
Predicted vs. observed abundance (A) of gypsy moth males in Virginia.

| Year | Observed maximum daily catch ($M_{pd}$) | Observed flight period (weeks) | Predicted Range of Abundance (A)<br>$0.96M_{pd}F < A < 2.87M_{pd}F$ | Observed abundance (A) |
| --- | --- | --- | --- | --- |
| 2013 | 13.5 | 6 | $77.76 < A < 232.47$ | 157.5 |
| 2016 | 15 | 6 | $86.4 < A < 258.3$ | 119 |

## Discussion

In this paper, we have established a predictive linear relationship between insect catch *M*_*pw*_ during the week of peak abundance, the length of seasonal flight period, *F* (number of weeks) and season-long cumulative catch *A* ≈ 0.41*M*_*pw*_*F*. This relationship was derived without any fitting to experiment, based on very general assumptions, which likely explains its remarkable accuracy and the fact that it works across the three tested species with very different biology, behavior, and trapping methods: *Lymantria dispar* (Lepidoptera: Lymantriidae), *Culex pipiens* (Diptera: Culicidae) and *Aedes albopictus* (Diptera: Culicidae). Although the model has only been validated using three species of insects, we expect the relationship, Equation 1, between maximum weekly catches and season-long catches to hold true for many other species, as long as their emergence has a clear start and end points, a clear maximum, and as long the collection method is sensitive and faithfully reflects variation in abundance. A quantitative criterion for the model applicability is also provided. The expectation that the model will work for many species is based on the generality of the arguments used in the derivation, absence of fitting to a specific species or trap type, and the already established agreement with experiment across three species.

For a theoretical model to work for a very broad variety of animal species, it must be based on very general principles that transgress specific biology of individual species. A good existing example is allometric laws in biology, which relate various biological characteristics to animal body mass. Originally, some of these laws were established purely empirically for a handful of species, then rigorously derived [41] based on general principles of energy conservation and distribution networks, which extended their applicability to essentially any species.

The data collected on gypsy moth phenology over the past 16 years allowed us to conduct a comprehensive analysis and relate season-long and weekly trap catches and flight duration to the daily trap catches. The data collectively indicate a significant variability in flight duration as estimated using pheromone-baited traps, which in turn causes significant variability in peak trap catches in populations with the same density as measured by male moth catches in pheromone-baited traps. To account for this observed variability, the model provides a range for a daily peak value, to allow researches and managers to estimate best and worst case scenarios, predict efficacy of control tactic, and make decisions to ensure optimal results. In addition to the immediate application in gypsy moth management programs, this model may be utilized to predict mating success of gypsy moth females and likelihood of persistence of isolated low-density populations [6, 42, 43]. Daily trap catch data collected over the entire activity season are not easy to obtain, however, should this data become available for species other than gypsy moth, it would be interesting to evaluate this model for these species as well.

Two interrelated scenarios of the potential model use in practical applications are exemplified below:

### Example Usage Scenario I

Equation 1, A ≈ 0.41 *M*_*pw*_*F*, can be used to estimate the hard-to-measure peak abundance Mpw from the other two population parameters, *A* and *F*, which themselves do not require intense monitoring. Thus, data collected as part of monitoring in the pest management program can be analyzed using this formula and appropriate method of control can be chosen based on the maximum daily value. This is important in large-scale control/monitoring programs, in which a method of control depends on the population density. For example, currently, in the Slow The Spread of the gypsy moth program (STS), the low dosage (6 g AI/acre) of mating disruptant is recommended for use in low-density gypsy moth populations, in which trap catches do not exceed 30 males/trap/season [7]. Ability to directly relate weekly trap catches from experimental plots to the season-long trap catches used by STS decision algorithm [44, 45] may allow us to improve the method of mating disruption used against gypsy moth by accurately estimating population densities during peak flight and using more appropriate control measures.

### Example Usage Scenario II

Equation 3, 0.96*M*_*pd*_*F* < A < 2.87*M*_*pd*_*F*, can be used to estimate the unknown season-long trap catch for a population based on the maximum daily peak abundance.

This is useful for model (artificially created) populations used in research studies, in which maximum daily values are available by experimental design, while the season-long trap catches are not available. In many mark – release – recapture studies, researchers are not trying to simulate flight, they simply repeatedly release similar numbers of insects to assess treatment efficacy. Therefore, the total of all captured insects in an experimental plot cannot be interpreted as the true season-long catch, which, therefore, would be unknown. Instead, each trap catch can be treated as the maximum daily catch during peak flight and the season-long trap catch can be estimated using the proposed formula. This approach allows to estimate the population density for which the treatment efficacy is assessed, to better interpret research results and to appropriately apply them in the management programs [46].

## Acknowledgements

We thank Slow the Spread of the gypsy moth Program Technical committee for helpful discussions and support for the research that lead to this analysis; Andrea Hickman, Samuel Newcomer, and Michael Merz for field assistance; Timothy McGonegal for providing mosquito trapping data. We also thank Andrew Liebhold (USDA Forest Service) for his valuable comments on an earlier version of the manuscript.

